# Online sensory feedback during active search improves tactile localization

**DOI:** 10.1101/590539

**Authors:** Xaver Fuchs, Dirk U. Wulff, Tobias Heed

## Abstract

Many natural behaviors involve closed feedback loops in which ongoing sensory input refines motor behavior. Previous research on tactile localization, however, has implemented localization as open-loop behavior. For instance, participants indicate a touched position on a silhouette shape of the body or on an occluding board mounted above the hand. Such studies have suggested that humans often make large errors when localizing touch on the skin, or that “perceptual body representations” are distorted. However, by artificially preventing tactile feedback from the target body area, the natural action-perception loop is interrupted. Therefore, these localization approaches may underestimate individuals’ localization ability and draw erroneous conclusions about the role and precision of body representations. Here, we tested tactile localization in a natural setting, in which participants first received brief touches on their left forearm and then searched for the target location by moving the right index finger across the skin. Tactile search reduced localization error when the searching finger was far from, but not when it was near the target, resulting in a remaining error of 1-2 cm. Error reduction was absent when participants searched on an acrylic barrier mounted above the arm, suggesting that availability of tactile feedback on the target arm but not proprioceptive and motor signals of the searching arm determined precision, thus confirming the pivotal role of closed-loop sensory feedback for tactile localization. We suggest that actively produced online tactile feedback routinely refines coarse spatial body representations, similar to the refinement of sparse spatial representations in visual memory through consecutive saccades.

## Introduction

Localizing touch on the skin is a fundamental function of the tactile system. Yet, humans often misjudge tactile location by up to several centimeters, even in seemingly simple tasks such as pointing towards the touched body part (1–3). Different factors may affect localization accuracy. On many body parts, receptor density is surprisingly low, limiting the acuity with which stimulus location on the skin can be detected (4). Furthermore, touch occurs on the two-dimensional sheet of the skin, and must be combined with posture to derive a location in space, before a goal-directed motor response towards the touch can be executed; these sensorimotor transformations potentially introduce further error (5–8). Lastly, localization error can result from motor error of the acting limb because motor execution is not performed with perfect reliability, as is evident from the observation that reaching endpoint varies across repeated trials even if the movement target is stationary (9, 10).

Researchers have investigated tactile localization with various experimental paradigms, mostly on the upper limbs. Some studies have asked individuals to indicate touch locations on a silhouette shape of the body (11, 12) or to assign touch location to a grid drawn on the limb (4). Others have required reaching or pointing movements without touching the target limb (13), or with the tactile target hidden under an occluding board (14) or touch screen (15).

One important aspect shared by all these methods is that participants, on purpose, do not receive tactile feedback about their localization accuracy from the target region. Thus, all above-mentioned methods are open-loop. In contrast, real-life situations often involve touching one’s own skin when localizing the stimulus and, thus, allow closed-loop control, underlining the active nature of seemingly perceptual functions. The experimental open-loop constraint is typically based on a purposeful decision aimed to study “pure” tactile localization or “perceptual” body representations and avoid potentially confounding influence of additional tactile information produced during localization. As one example, open-loop responses of tactile localization on the hand do not map onto the actual dimensions of the hand, suggesting that the brain represents the hand as shorter and wider than it actually is (16, 17). Yet, others have criticized these conclusions, arguing that distortions may be due to location estimates being referred to previous estimates, that is, that they may originate from domain-general bias induced by the required behavioral response rather than from true representational distortion (18).

Beyond the context of tactile localization, the notion that perception is an active process that encompasses an interplay between sensory and motor processes is a widely accepted idea (19). For instance, tactile object recognition critically depends on manipulating the object with the hands, and moving one’s fingers across it (20). Furthermore, spatial perception in touch can be anchored to the timing of active movement, as demonstrated by neural coding of touch during whisking in rats (21), as well as systematic localization errors that arise when humans indicate the location of touch that occurred during arm movements (22). In vision, saccades into the periphery often slightly miss their target, and are then corrected by secondary saccades, both in the laboratory (23, 24) and naturalistic environments (25). Note, that the visual target falls on the low-resolution periphery of the eye before the first saccade. The spatial location of the target can, therefore, not be precisely determined. The first saccade then moves the visual target into the high-resolution fovea, allowing online assessment of exact target location and, hence, correction of remaining error. We hypothesized that a similar closed-loop mechanism may support tactile localization. In particular, tactile localization may be coarse, with online tactile feedback during search providing higher-resolution tactile-spatial information. Imagine scratching your arm: often, we simply direct our scratching hand grossly towards the itch. Then, we scratch across the itching arm until we find the location from where the itch originates.

Notably, two distinct mechanisms may underlie such behavior. A closed-loop account posits that searching movements should be systematically directed towards the target location, that is, the distance between the acting hand and the tactile target should reduce continually. An alternative possibility, however, is that the hand searches randomly, rather than goal-directed, so that success in search would depend on hitting the target by chance during extended movements that are unrelated to target location, rather than on a closed-loop sensorimotor control mechanism.

## Results and discussion

In experiment 1, fifty-eight healthy human participants received brief tactile stimuli, applied with a wooden stick, in one of three locations on the left dorsal forearm: a proximal position near the elbow, a distal position near the wrist, and a medial position between them. In two separate blocks, participants held their arm orthogonal or parallel to the torso (straight vs. angled posture, Fig. 1a,b). Because low-level skin maps are thought to be anatomically organized (26), common external-spatial error across the two postures would indicate motor error of the pointing hand rather than perceptual localization error on the target arm. Participants reached, with their eyes closed, using their right index finger to the left forearm, touched down, and then moved their finger across the skin until they felt to have reached the target. Stimulus and finger locations were recorded from above with a camera. Finger movement trajectories on the skin were extracted from video and the coordinates were transformed into units of percent forearm length (see Methods).

**Figure 1:**
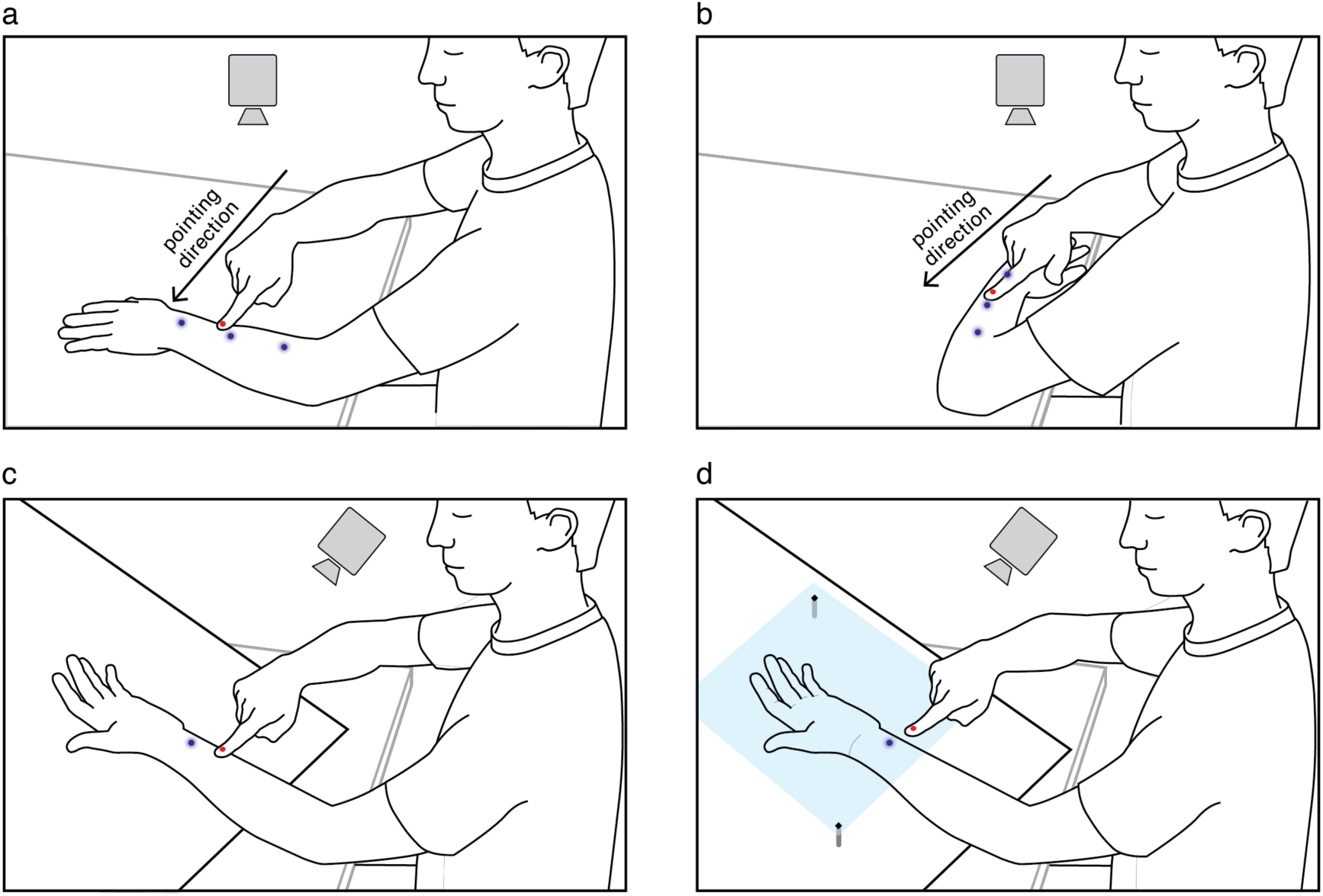
Illustration of Experiments 1 and 2. (a, b) In Experiment 1, participants received brief touches, applied with a hand-held wooden stick, in one of three target areas of the dorsal forearm (blue points). Subsequently, they localized the touch with their right hand’s index finger. Two posture orientations, straight (a) and angled (b), were used to separate perceptual from motor localization error: The pointing hand approaches the target arm from the right (from the participant’s view) in both posture orientations. Hence, motor error (e.g. overshoot) would affect different anatomical on the target arm for the two postures. (c, d) In Experiment 2, touch was applied to the left ventral forearm. The participants localized the stimuli by either moving the finger on the skin (c), as in Experiment 1, or by moving the finger on an acrylic glass barrier (d), preventing tactile feedback from the skin of the target arm.

### Constant and variable localization error

We first evaluated the constant error in proximodistal (from the elbow to the wrist) and mediolateral direction (from the inside to the outside of the arm). Constant errors express systematic localization biases or distortions in body representations (14, 16). We computed constant errors for initial and final localization, that is, at touch-down of the finger versus after tactile search on the skin (Fig. 2). Initial and final constant error were computed across trials as the intra-subject mean distance between the target and the initial and final finger location, respectively.

**Figure 2:**
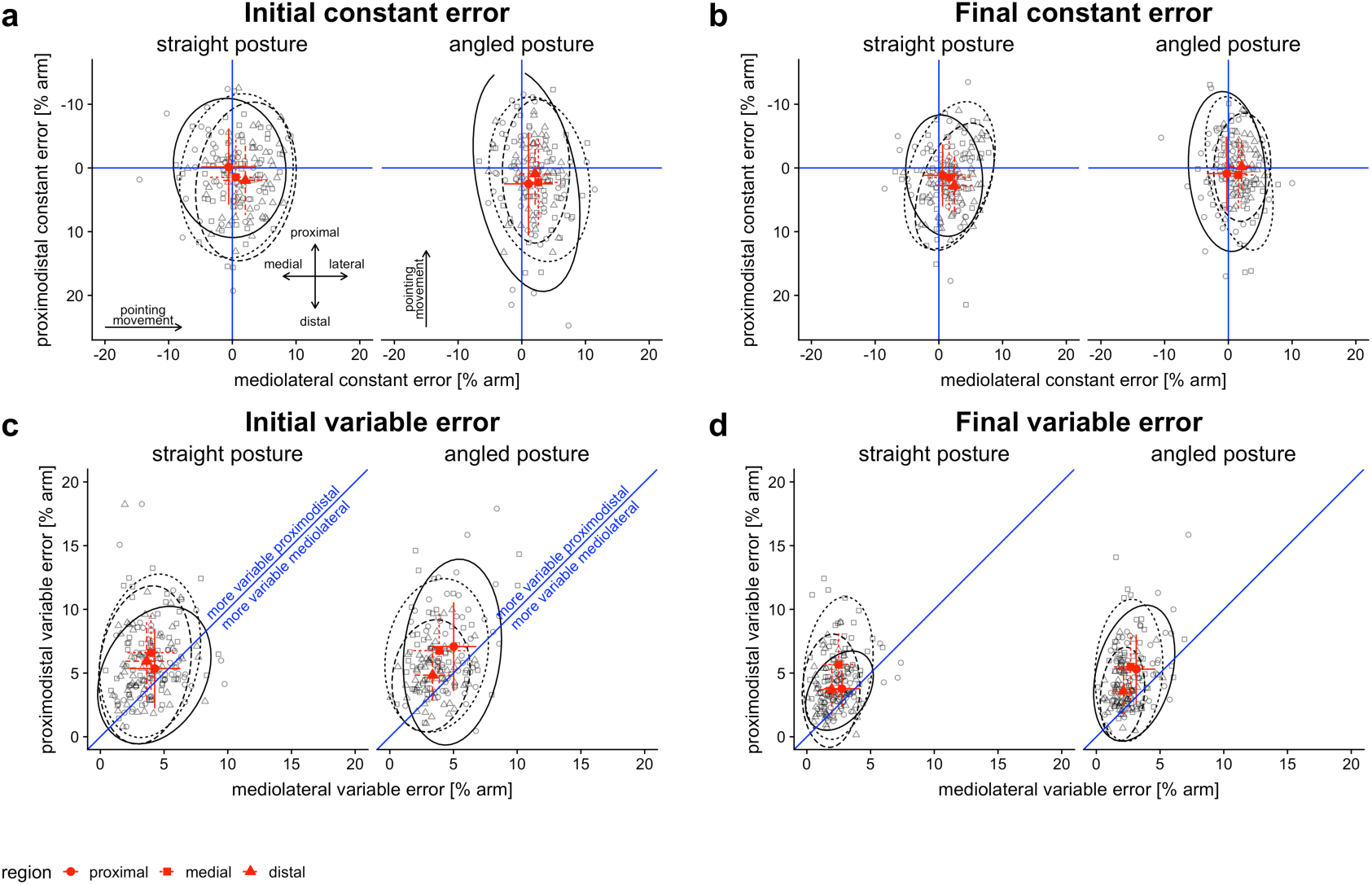
Constant and variable error in Experiment 1. (a, b) Constant error in mediolateral and proximodistal direction for the straight and the angled posture for initial localizations. Error was computed as the intra-subject mean localization error over trials, coded as distance from the target, represented by the crossing point of the blue lines. Single participants’ error is shown as black points separately for the three target regions (proximal, medial and distal target). The red symbols with crosses indicate group means for the target regions with standard deviations as error bars. The ellipses represent 95 % confidence. The lower-right quadrant of panel a and b indicates bias in distal and lateral direction. (a) At finger touch-down, before search, (b) after search. (c, d) Variable error was computed as the intra-subject standard deviation of localization error over trials and therefore express participants’ precision. Values that fall into the area above the blue diagonal line indicate larger variable errors in proximodistal compared to mediolateral direction. (c) At finger touch-down, before search, (d) after search.

For the final localization, we observed constant errors in the distal, i.e., towards the hand (2% arm length, p < 0.001), and in the lateral direction, i.e., towards the outside of the arm (1% arm length, p < 0.001). Constant error magnitude differed between target regions (Fig. 2 and Table S2 in the Supplementary Material). There was no overall difference between arm postures (F(1, 1274) = 0.37, p = 0.54), suggesting that constant error was not a result of pointing movement direction. However, differences between target regions depended on posture (F(2, 1239) = 5.76, p = 0.003; see Table S1 in the Supplementary Material for details). Overall, constant error was of comparable magnitude for initial finger location, ruling out a contribution of search on the skin to localization bias (F(1, 1239) = 0.26, p = 0.610). Constant error was also statistically significant for each posture both before and after search (see Table S4 in the Supplementary Material). However, while not changing the general direction of the errors, distal and lateral bias slightly diminished after search in the angled but increased in the straight posture (F(1, 1239) = 13.99, p < 0.001). The present distal localization biases are in line with previous reports based on various measurement methods, suggesting distorted body representations (14, 16).

Whereas tactile search did not compensate for perceptual bias, it did reduce the absolute distance from the target (F(1, 1240) = 19.2, p < 0.001). Error reduction was evident in both distal and lateral direction (Fig. 2a vs. 2b; Fig. 4a; Table S6 in the Supplementary Material). Absolute error was larger in proximodistal than in mediolateral direction (4.5% vs. 2.9% arm length, p < 0.001; Fig. 2a,b).

The reduction in absolute error through tactile search was accompanied by a reduction in variable error (F(1, 1223) = 174.3, p < 0.001). Variable error describes the variability in localization performance (unsystematic error) and is computed as the intra-subject standard deviation across trials. Overall, variable error was larger, that is, localization performance was more variable, in proximodistal than in mediolateral direction (5.3% vs. 3.3% arm length, p < 0.001, Fig. 2c,d). These effects were evident across all conditions, although error was slightly larger in the angled than the straight posture, and it differed between the target regions (Fig., 2c,d and Table S8-S10 in the Supplementary Material).

The reduction in variable error, together with the findings of invariable constant error across search and invariable improvements across postures, implies that localizations gravitated towards the target location and, hence, search reduced localization error from different directions.

Because tactile search reduced localization error in all directions, we conducted further analyses on the Euclidian distance to the target, thus collapsing error into a single, direction-independent, measure. Tactile search improved final localization in 68-72% of trials for the different conditions, confirming that error reduction was a general effect and did not, for instance, depend on just a few trials with very large corrections (Fig. 3a). Similarly, the variance across trials in Euclidian distance was lower for final than for initial localization in 57-81% of the participants for the different target regions (Fig. 3b).

**Figure 3:**
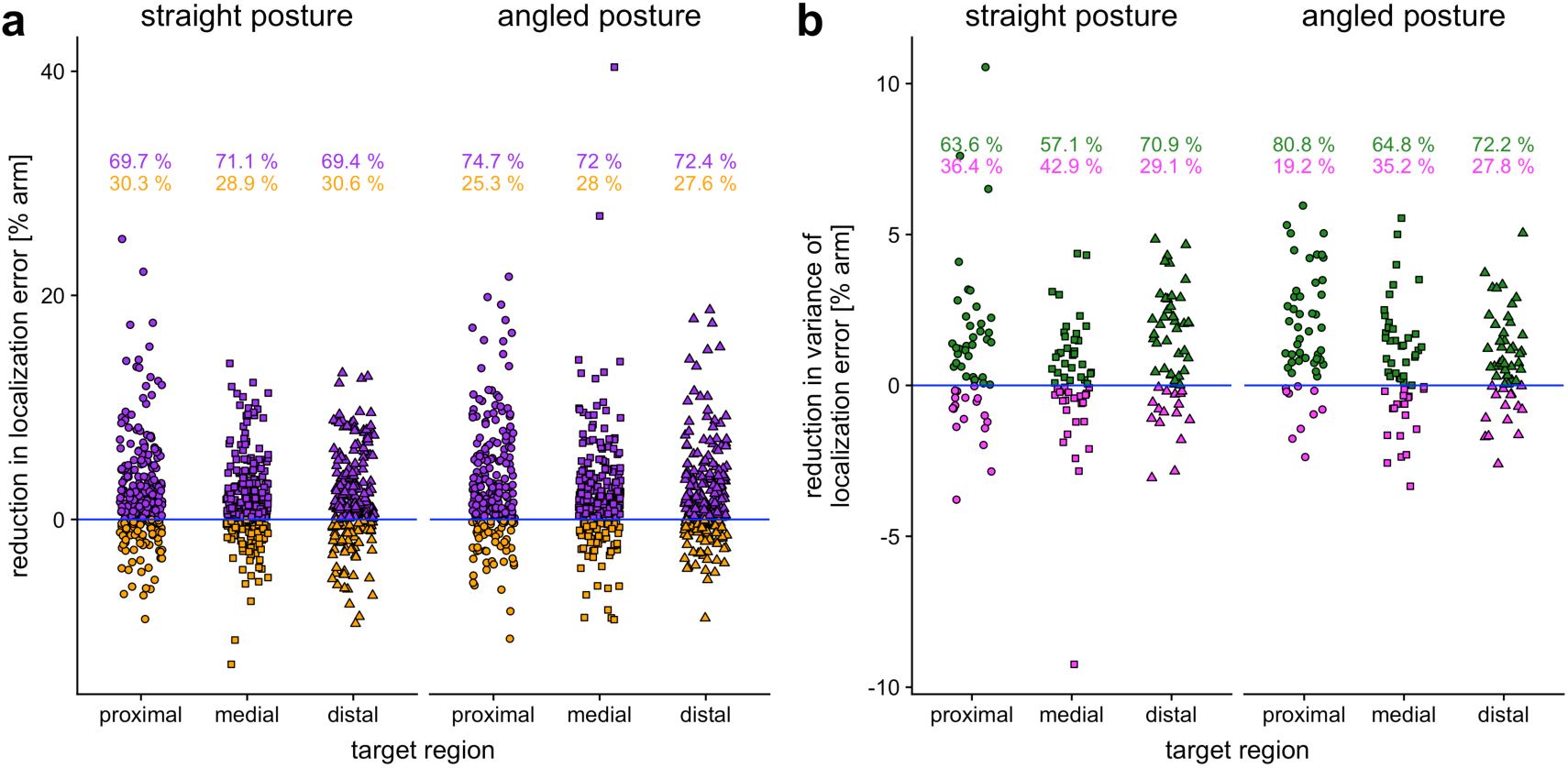
Reduction of direction-independent tactile localization error (computed as Euclidian distance from the target) through search in Experiment 1. (a) Intra-subject difference values between localization error before vs. after tactile search. Each point represents one trial. (b) Difference values between intra-subject variation (computed as standard deviations) of localization error before search vs. after search. Positive values indicate smaller variation after search than before, indicating that, through search, localizations gravitate towards their mean value.

### The beginning, but not the end, of the search trajectory reduces localization error

So far, we have established that tactile search reduced localization error. To better characterize the observed error reduction, we analyzed how it developed across the search trajectory and whether it depended on the magnitude of the initial localization error. To this end, we first split search trajectories into 20 segments of equal spatial length to control for difference in movement time and speed (see Methods). Note that this analysis step fully reduces trajectories to their spatial features (in this case to segments representing steps of 5% of the trajectory length). Accordingly, the recoded trajectories are not influenced by the fact that original movement trajectories at constant sample rate usually have many data points at the beginning and the end due to the lower movement speed.

We analyzed error reduction across the 20 segments separately for trials with initially-large, initially-medium, and initially-small localization error at touch-down. Localization error diminished non-linearly across segments: reduction was greatest during the first search segments, and absent in late segments (Fig. 4a,b). However, this effect critically depended on initial localization error: The linear term of a polynomial regression was strongly negative for initially-large-error trials, indicating continuous improvement in localization over the course of the trial (b = −7.0, p < 0.001). In contrast, it was near zero but slightly positive for initially-small-error trials, indicating even slight deterioration of localization over segments (b = 0.83, p < 0.001). Moreover, the quadratic term was strongly positive for the initially-large-error trials, indicating that localization improvement was large at the beginning, and absent towards the end of search (b = 2.4, p < 0.001). In contrast, the quadratic term was small and negative for initially-small-error trials, indicating that localization deteriorated at the beginning, and remained unchanged at the end of search (b = −0.92, p < 0.001). Thus, tactile search significantly reduced localization error at the beginning of search only when participants had set their finger down relatively far from the target. Independent of initial distance to the target, search was ineffective towards the end of search. For trials in which initial localization error was of medium size, both the linear and the quadratic terms fell in between those of the large and small error trials, suggesting a graded effect of initial localization error. The effect that localization error reduced for initial large, but not for small localization error, was present for both arm postures and all three target regions (Fig 4b and Table S13 in the Supplementary Material).

**Figure 4:**
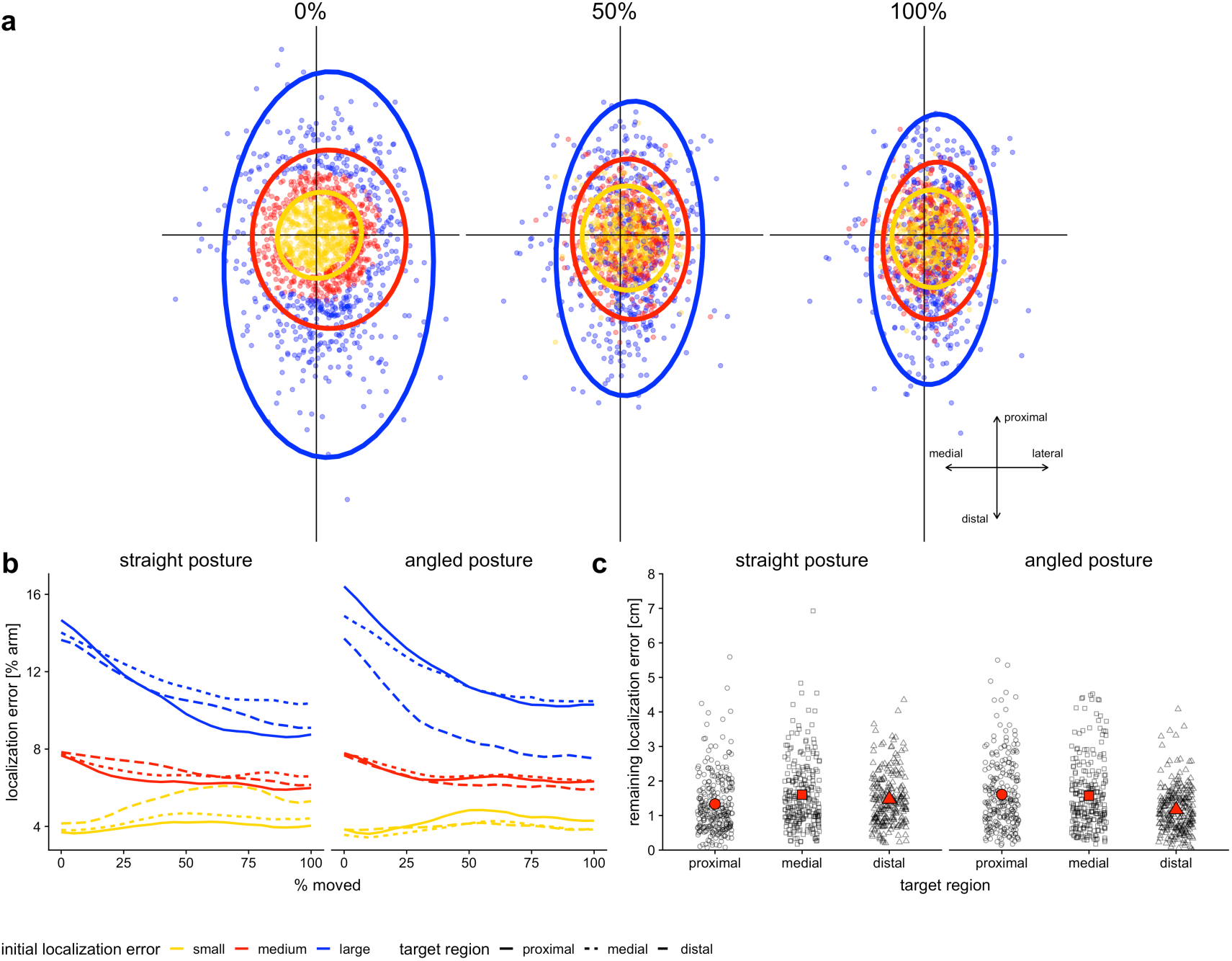
(a) Distribution of single trial localization error in Experiment 1, pooled across arm postures and target regions. Localization error is defined as the distance of the pointing finger from target position, represented by crossing lines in the middle of the plots. Trials are divided into three groups according to initial distance from the target (blue: far, red: medium, yellow: close) at touch-down. Ellipses represent 95% confidence for the respective, color-coded trial group. From left to right, at touch-down, after 50% of the travelled distance, and at end of search. An animated version of this figure is available online as Supplementary Material. (b) Change of direction-independent localization error (defined as the Euclidian distance from the target) over the course of tactile search in dependence of initial localization error. (c) Remaining direction-independent localization error, in centimeter units, at the end of tactile search.

Results so far suggest that search was based on a closed-loop strategy that consistently reduced error when it was large. However, the absence of localization error reduction in initially-small error trials and the later segments of initially-large-error trials implies that error reduction is confined to a region that is distant from the target. One possible cause of this limit may be the large size of tactile receptive fields on the arm (4). Consistent with this interpretation, we found the average final error (Fig. 4d) to be 1-2 cm, which roughly matches existing estimates of the two-point discrimination threshold on the arm (4).

### Error reduction by search is due to tactile information on the target limb

The results presented above suggest that participants rely on tactile feedback around the target for error reduction during tactile search. One alternative interpretation, however, is that search is driven by motor correction and proprioceptive feedback of the acting, searching arm. Under this notion, participants used a precise tactile target location estimate throughout, but had to gradually direct their searching finger to that location due to imprecision of the motor system. Experiment 2 scrutinized this alternative interpretation (see Fig. 1c,d). Sixteen new participants localized tactile stimuli on the left, extended, ventral forearm with their right index finger either searching on the skin as in Experiment 1, or on an acrylic glass barrier directly above the arm (14). The barrier prevented tactile feedback of the target arm.

Search on the skin reduced localization error, especially in the first half of search (linear term: b = −4.4, p < 0.001; quadratic term: b = 1.7, p = 0.004; Fig. 5a,c). Again, error reduction depended on initial localization error and was strongest in trials with initially large localization error, whereas it was weak and reversed when initial localization error was small (F(2, 6072) = 73.6, p < 0.001; see Table S17 and S18 in the Supplementary Material). Thus, Experiment 2 replicated the results from Experiment 1, but for a different region of the skin. Yet, in line with previous studies (16), we did not observe any distal constant error (all p > 0.05; see Table S15 in the Supplementary Material) on the ventral as opposed to on the dorsal forearm.

**Figure 5:**
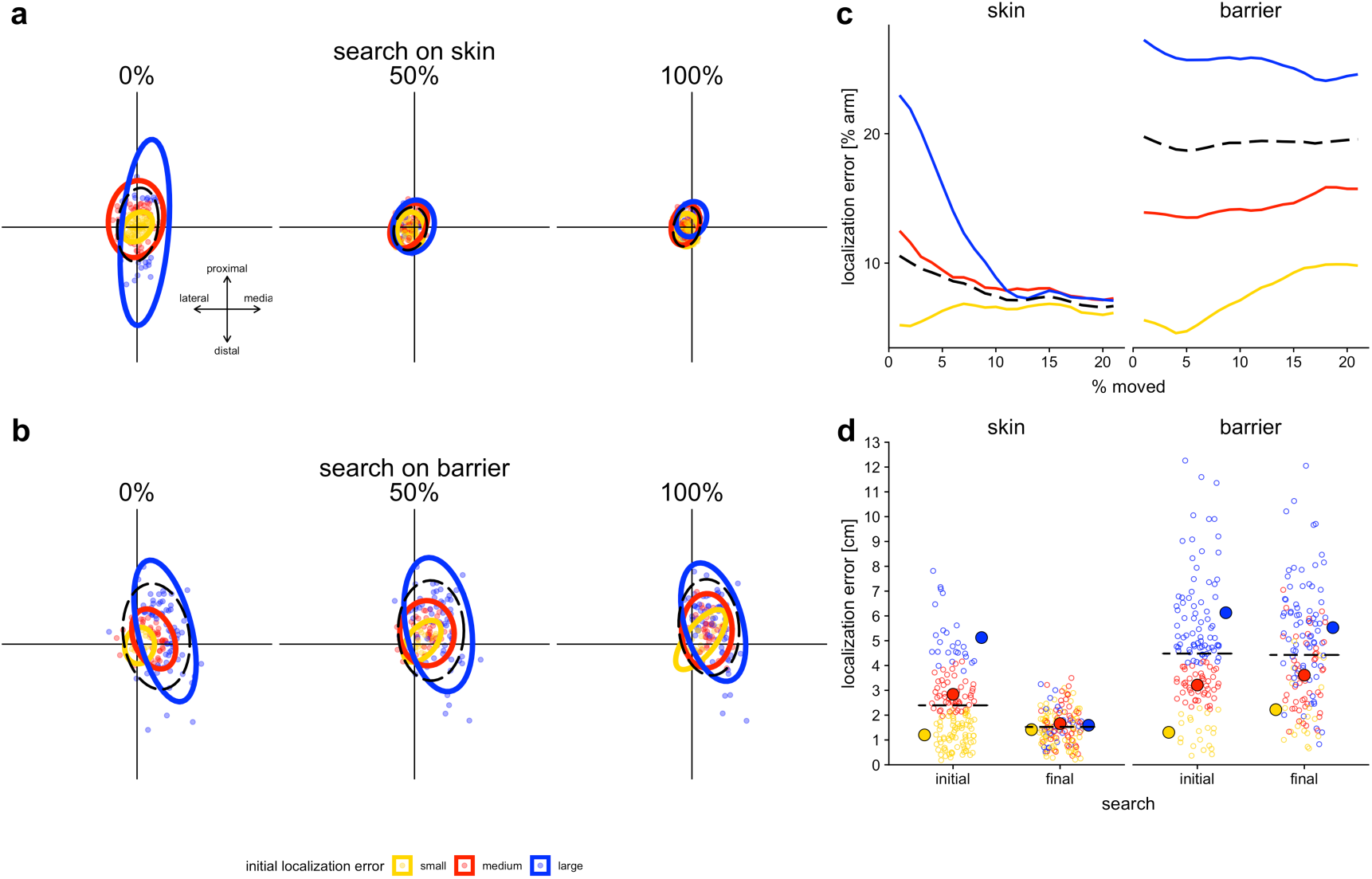
Results of Experiment 2. (a) Localization error distribution of single trials for search with tactile feedback, comparable to Experiment 1 (compare to Fig. 3a) (b) Localization error distributions for search on an acrylic barrier, that is, without tactile feedback on the target arm. An animated version of (a) and (b) is available online in the Supplementary Material. (c) Change of localization error over the course of search depending on initial distance from the target for tactile feedback and acrylic barrier conditions. (d) Remaining distance from the target after search. Colors code subgroups with different initial distance from the target; the dotted lines indicate the average of all trial types collapsed.

Critically, we observed significant constant error in the barrier condition in proximal (4.5% arm length before search, 9% after search, both p < 0.001) and in medial direction (12% arm length before search, 11% after search, both p < 0.001; Fig. 5b and Table S15 in the Supplementary Material). Thus, prohibiting closed-loop feedback by an occluding barrier introduced systematic constant error that would classically be interpreted as a perceptual or representational distortion, demonstrating that the choice of experimental localization response can affect the qualitative conclusions drawn by the respective localization error. Moreover, localization on the acrylic barrier produced significantly larger variable localization error than search on the skin (F(1, 233) = 131.9, p < 0.001, Fig. 5c). In fact, the average localization error for barrier search was 4.5 cm before, and 4.4 cm after search, and, thus, nearly three times larger than the remaining error of 1.5 cm for skin search (Fig. 5d). Neither the linear nor the quadratic term were statistically significant in the 20-segment model for acrylic search (linear term: b = 0.53, p = 0.453; quadratic term: b = 0.36, p = 0.607; Fig. 4b,c and Table S18 in the Supplementary Material). In relative terms, these results reflect an improvement of merely 1% for search on the barrier and, on average, 36% for search on the skin (initially-large/medium/small-error: - 17/41%/69%). We conclude that closed-loop tactile information on the target region, but not proprioceptive information of the acting arm, considerably improved tactile localization accuracy and precision.

## General discussion

Human perception of touch location is subject to multiple biases and illusions (14, 27), and these distortions are likely related to the shape of receptive fields and low tactile receptor density of many body regions (4, 28). However, virtually all methods that are currently employed in tactile localization research and body representation deliberately prevent tactile feedback of the target region, presumably to focus on “pure” perceptual processing. The present results indicate that this strategy not only risks grossly underestimating human participants’ tactile localization ability in more natural situations, and may introduce systematic distortions that reflect methodological limitations rather than cognitive principles. Thus, attempts to isolate perceptual or representational aspects of tactile-spatial processing must be carefully and thoroughly pitted against experimental paradigms that engage complete perception-action loops if unwarranted conclusions are to be excluded with confidence. Our findings emphasize that tactile localization is an active search process that relies on closed-loop, tactile feedback and challenge the validity of approaches that prevent tactile feedback in studies on tactile localization.

In the visual domain, it has been suggested that spatial representations are sparse for the purpose of reducing memory load and energy consumption, but are effortlessly updated by sensory input generated through saccades (29, 30) – effectively relying on the “world as an outside memory” (29, 31). Both in the laboratory (23, 24) and in naturalistic environments (25), initial saccades regularly miss peripheral targets, evoking corrective saccades upon detection of the remaining error. By analogy, the present results may reflect a purposeful strategy to maintain globally sparse body representations that, when required, can be locally enhanced online by means of active search. Such a strategy appears both efficient in terms of computational costs and biologically plausible. In this view, the sparsity of representations is not accompanied by any major functional disadvantages as they can efficiently be remedied through purposeful behavior.

It is remarkable that search was entirely ineffective, and even slightly detrimental, when initial target error was small, and that search continued after initial error reduction, without any further error-reducing effect. This search behavior appears consistent with random search that attempts to detect the remembered location through chance movement rather than through goal-directed, closed-loop error correction. However, given that initial search is efficient with respect error reduction, a more likely possibility is that participants adjust movement close to the target based on random fluctuations of tactile sensation, and end the search after they have been unable to further reduce their average error after some time has elapsed, akin to decision theories that posit that urgency signals are instrumental in forming sensory decisions (32). Notably, although a remaining error of 1-2 cm may appear large, this accuracy of localization is probably sufficient in everyday life. With only very few exceptions, objects touching our skin cover an area, and not just a point, of the skin. Furthermore, we reduce finger location to a point for analysis, but in truth the finger is about 1.5 cm wide even at its tip; as is the case for any study speaking of limb position as a single coordinate, it is therefore questionable how exact the spatial resolution of body part position can be. Thus, although our results shed light on the mechanism employed in tactile search, in practical terms, the final error we determined in our study is likely to be unnoticeable outside the laboratory.

To summarize, error reduction through search was entirely due to tactile feedback from the target arm and confined to a distance of 1-2 cm around the target, roughly matching the two-point discrimination threshold. Tactile localization is a closed-loop, active process, in which online sensory input reduces localization error. Therefore, open-loop procedures interrupt the localization process; relying on localization while prohibiting tactile feedback of the target region underestimates localization ability and may introduce systematic bias.

## Methods

We conducted two experiments. For Experiment 1, our aim was to assess pointing movements to tactile events in a large and heterogeneous sample, in a setting that was as natural as possible while allowing performance assessment. For this purpose, we recruited participants at a public science fair in Bielefeld. For Experiment 2, our aim was to explicitly control some factors which we had ignored in Experiment 1 in exchange for a more natural paradigm. Experiment 2 was, therefore, performed in the lab. The experiments were approved by the Ethics Committee of Bielefeld University, Bielefeld, Germany.

### Participants

#### Experiment 1

We recruited 58 participants for Experiment 1 (aged 14-73, mean = 40.5 years; 36 females, 21 males, in one case sex was not assessed). Fifty participants were right handed, 5 were bidexterous and 3 were left handed, according to self-report. Participants gave informed, verbal consent, and did not receive monetary reward.

#### Experiment 2

Sixteen participants (aged 19-34, mean = 25.4 years; 8 females) participated either without compensation or for course credit. All participants were right-handed, according to the Edinburgh Handedness Inventory (33), and reported to be free of neurological or psychological disorder or any other condition that might affect sensory or cognitive function. Participants gave written, informed consent.

### Procedures

#### General procedure

In both experiments, participants received, in each trial, a brief touch on their left forearm, and subsequently indicated tactile location by pointing with their right index finger. In contrast to previous research, participants were explicitly allowed, and moreover encouraged, to move their index finger across the target arm’s skin during localization. We measured the length of the target forearm, from the crook of the elbow to the wrist, using a measuring tape. Then, participants performed a few practice trials until they indicated that they had understood the procedure.

##### Experiment 1

Participants closed their eyes during trials. Stimulation was applied to the dorsal forearm. Participants initiate the pointing movement when the tactile stimulus had been released from the skin. When participants were done with their search, they kept their finger stationary in the final position for a few seconds before they returned to the starting position 39 cm to the right of the target arm. The experiment took about 15 minutes.

##### Experiment 2

Participants were blindfolded during the experiment. Stimulation was applied to the ventral forearm. We chose a different stimulation site than in Experiment 1 to test whether our results would generalize to different surfaces. Localization was paced in order to keep constant the time that passed between stimulus and localization across trials and conditions. Experiment 2 involved a condition in which participants searched on a barrier above their arm, rather than on the arm itself. Because this barrier was placed as close to the arm as possible, it had to be removed for stimulation, and then be replaced before search. A timer instructed participants to refrain from initiating the search until 3 s following stimulation, resulting in identical trial timing for skin and barrier conditions. The experiment took about 30 minutes.

#### Tactile stimuli

Tactile stimulation was applied by an experimenter for approximately 1 s.

##### Experiment 1

Stimuli were applied via a hand-held, wooden stick of approximately 35 cm length. The stick’s edges were blunt and evoked a light touch sensation. We applied a force that caused a visible dent and slight whitening of the skin in the target region.

##### Experiment 2

To standardize touch force, we applied tactile stimulation with a von-Frey filament (Marstock Nervtest, Schriesheim, Germany) with 256 mN force.

#### Experimental conditions

##### Experiment 1

We manipulated two experimental factors. First, *target arm posture* varied between a straight and an angled posture, the latter with the elbow joint at approximately 90° (see Fig. 1). Posture was varied blockwise, with the order balanced across participants. Second, target region between three different areas along the forearm: a proximal position near the elbow; a medial position half-way between elbow crook and wrist; and a distal position near the wrist. Targets were always centered with respect to the mediolateral axis of the arm, that is, stimuli were always applied on top of the arm. Within the three defined regions, we slightly jittered location between trials along the proximodistal axis to avoid adaption or practice effects. Within the posture blocks, each region was touched five times in pseudorandomized order.

##### Experiment 2

We varied whether participants received tactile feedback from the target region. Participants either localized stimuli by searching on their skin versus on the acrylic barrier. For both conditions, participants positioned their left arm in a wooden construction (60 × 39.2 cm, see Fig. 1), with the ventral forearm surface turned towards the face. Their elbow rested comfortably on the table surface and the construction maintained the elbow at an angle of 120°. We used two target locations 4 cm proximal of the center of the wrist that were 1 cm apart in mediolateral direction. For precise stimulation, we marked the targets on the arm with a pen. Multiple dots were drawn in a 10×10 cm area around the target to obscure the true location to participants during preparation. During the experiment proper, participants did not see these locations due to the blindfold. Each target was repeated five times. An acrylic barrier of 15 × 15 cm was placed above the forearm. It could be attached at a flexible distance, so that it was positioned just above the arm without touching it. The barrier was removed for stimulation, and then replaced before localization.

#### Acquisition of stimulus positions and pointing movements

We recorded stimulus presentation and participants’ finger movement during localization with a digital camera (Intel RealSense, Intel, Santa Clara, California, USA) at a spatial resolution of 1280 × 720 pixels and a framerate of 30 frames/s. The camera was mounted above the table, aligned with the table surface in Experiment 1, and with the wooden construction in Experiment 2, to yield undistorted image data. The barrier in Experiment 2 was transparent and allowed recording the arm during search. A 0.8 cm diameter round, bright orange sticker on the participant’s right index finger’s nail indicated finger position. The camera was controlled using custom programming code in PsychoPy (v1.85.2, www.psychopy.org) running within the free Ubuntu operating system (v16.04, www.ubuntu.com) on a Laptop computer (Dell Latitude, Dell, Round Rock, Texas, USA).

#### Extraction of positions from image data

We analyzed videos with Matlab (v2016a, www.mathworks.com). First, we marked, for each trial, four anatomical coordinates of the forearm on the image, namely the inside and outside edges of the elbow and of the wrist. These four coordinates defined a trapezoid shape that approximately covered the area of the forearm. Second, we found the frame (i.e., timepoint in the video) in which the tactile stimulus was presented on the skin. We used the frame in which skin indentation was largest and marked stimulus location on the image. Third, we found the frame at which the participant’s finger first touched the target arm or barrier. Fourth, we found the frame in which the participant lifted off from the target arm. We then extracted the center position of the orange dot that marked index finger position for all frames between touchdown and liftoff. Note that the time passing before and after movement on the skin, when the participant’s finger stayed idle, is irrelevant for the statistical analyses of movement trajectories because we analyzed trajectories with respect to space, independent of time (see section on data processing).

#### Data loss and exclusion

##### Experiment 1

Five of originally 63 data sets were excluded due to technical problems. For the remaining 58 participants within the analyzed dataset, at least one trial was missing for 48 (85%) cases, with an overall trial loss of 15%. The analysis we present is based on 1480 trials. In one participant the information about arm length was missing and the value was substituted by the sample average.

##### Experiment 2

We excluded 4 of originally 20 tested participants from the analysis due to technical problems during acquisition. For the 16 remaining participants, at least one trial was missing in 12 (75%) cases, with overall data loss of 9%. The presented analysis is based on 555 trials.

### Statistical Analysis

#### Data preprocessing

All data were analyzed with R (v3.5.1., https://cran.r-project.org/) (34). First, we aligned all data spatially to eliminate differences due to posture and individuals’ limb size. For each trial, the arm area was transformed to a mean group template based on the four anatomical coordinates of the elbow and wrist using the “Morpho” package for R (35). For each single trial, arm area, tactile location, and the search trajectory were transformed into the group template space. Finally, we expressed all coordinates as percent of arm length (36), measured as the distance between the middle position between the two elbow coordinates and the middle between the two wrist coordinates of the template. As the outcome variable for statistical analysis, we expressed trajectories relative to target position, effectively coding trajectories as continuous localization error. For comparison with previous studies, we analyzed trajectories separately in proximodistal and in mediolateral direction. To reduce error into a single, direction-independent measure, we also expressed trajectories as the Euclidian distance between finger and target.

For the analysis of search progression, we “spatialized” trajectories using the R package “mousetrap” (37, 38). We recoded all trajectories into 20 segments between touch-down and lift-off. Following this step, each trajectory contained the same number of observations, and each segment coded 5% of the travelled distance, thus eliminating speed and time.

For the analysis of remaining error after search, we transformed the data into cm units by multiplying the percentage of arm length values with the participant-specific arm length in cm.

#### Statistical analysis

##### Experiment 1

We analyzed our data with linear mixed models with the “lme4” package for R (39). Linear mixed models account for differences in trial numbers between participants and allow for individually sized effects across participants. For analysis of constant and variable error in Experiment 1, we computed a random intercept model, including *posture* (straight and angled), *direction* (proximodistal and mediolateral), *target region* (proximal, medial, and distal), and *search* (initial and final localization) as fixed factors, including all possible interactions. We included participant as a random intercept effect, resulting in the formula, in lme4 notation: lmer(constError∼ posture*direction*region*search + (1|participant). To quantify the degree of bias, we computed estimated marginal means (EMMs) from the model using the “lmerTest” package (40). Because trajectories were coded as distance from the target, EMMs significantly different from zero indicate bias the respective direction. For constant error, we computed a model with the same factors, but with unsigned trajectory data as dependent measure. For analysis of variable error, we again computed a model but the same factors as before, but with unsigned trajectory data as dependent measure. To compare the different conditions. we contrasted their respective EMMs. We analyzed the progression of the search with a similar model that included fixed factors *posture* and *target region*. It used error in Euclidian space as dependent measure, rather than separating into proximodistal and mediolateral error, to reduce model complexity. The model expressed progression in the trial as a fixed factor *percent moved*, which consisted of twenty factor levels representing 5% spatial segments and used orthogonal polynomial contrasts of first and second order, i.e., a linear and a quadratic term. Furthermore, the model included *initial distance from the target* (initially-small, initially-medium, and initially-large error) as a fixed factor. This factor was defined by binning trials at percentile 33 and 66 according to the Euclidian distance from the target at touchdown. We ran submodels that included only the data of one factor level to break down significant interactions.

We considered effects to be statistically significant at p < 0.05. We corrected p-values for post-hoc tests for multiple comparisons using false discovery rate. *Experiment 2*. Analysis was analogous as for Experiment 1, but included as fixed factors *surface* (skin, barrier), *direction* (proximodistal and mediolateral), and *search* (initial and final localization).

## Data and Software Availability

All data and code used in this study are available on the website of the Open Science Framework (osf) and can be accessed at https://osf.io/v7hsj.

## Acknowledgments

This research/work was supported by the Emmy Noether grant of the German Research Foundation (DFG) to T.H. (He 6368/1-1/2/3) and by the DFG-funded Cluster of Excellence Cognitive Interaction Technology ‘CITEC’ (EXC 277) at Bielefeld University. We thank Marta Beauchamp, Joseph Gerges, Diana Kollenda, Mathilde Perchermeier, Vitalij Rozkov, Frederick Thiemer, and Lisa Viereck for support with participant recruitment, data acquisition and video processing, Conrad Alting for programming assistance, and Marta Beauchamp for contributing the drawings for Fig. 1.

## Author Contributions

Xaver Fuchs (XF) and Tobias Heed (TH) designed the experiments and analysis. XF conducted the experiments. XF and Dirk Wulff (DW) analyzed the data. XF prepared the figures. XF, DW, and TH wrote the manuscript. TH supervised the study.

