## Supplementary Material for "Online sensory feedback during active search improves tactile localization"

Table S13: Linear and quadratic terms for different initial error (small, medium, large), postures (straight vs. angled) and target regions (proximal, medial, distal) . 10

**2. Experiment 2** **11**

This document is a reproducible report, created with RMarkdown. Data and code to reproduce the analyses are provided in the accompanying repository on the website of the Open Science Framwork and can be accessed via the following link: https://osf.io/v7hsj

### 1. Experiment 1

#### 1.1. Statistical models for constant and variable error

##### 1.1.1. Linear mixed model for constant error

The following factors were included in the model:

- fixed factors:
  - posture (straight, angled)
  - direction (proximodistal, mediolateral)
  - target region (proximal, medial, distal)
  - search (initial final)
- random factors:
  - id (random intercept factor)

resulting formula in lme4 notation: `lmer(constError ~ posture*direction*region*search + (1|id))`

**Table S1: Anova table for constant error model**

Table resulting from R's anova() command.

| Term | Sum Sq | Mean Sq | NumDF | DenDF | F value | Pr(>F) |
|---|---|---|---|---|---|---|
| posture | 7.252 | 7.252 | 1 | 1275 | 0.368 | 0.5443 |
| direction | 0.183 | 0.183 | 1 | 1239 | 0.009 | 0.9232 |
| region | 316.415 | 158.208 | 2 | 1240 | 8.024 | <0.001 |
| search | 5.116 | 5.116 | 1 | 1239 | 0.259 | 0.6106 |
| posture:direction | 31.510 | 31.510 | 1 | 1239 | 1.598 | 0.2064 |
| posture:region | 227.066 | 113.533 | 2 | 1240 | 5.758 | 0.0032 |
| direction:region | 161.008 | 80.504 | 2 | 1239 | 4.083 | 0.0171 |
| posture:search | 275.797 | 275.797 | 1 | 1239 | 13.988 | <0.001 |
| direction:search | 9.924 | 9.924 | 1 | 1239 | 0.503 | 0.4782 |
| region:search | 4.706 | 2.353 | 2 | 1239 | 0.119 | 0.8875 |
| posture:direction:region | 97.117 | 48.559 | 2 | 1239 | 2.463 | 0.0856 |
| posture:direction:search | 2.667 | 2.667 | 1 | 1239 | 0.135 | 0.7131 |
| posture:region:search | 32.920 | 16.460 | 2 | 1239 | 0.835 | 0.4342 |
| direction:region:search | 3.455 | 1.728 | 2 | 1239 | 0.088 | 0.9161 |
| posture:direction:region:search | 20.883 | 10.441 | 2 | 1239 | 0.530 | 0.589 |

**Table S2: Estimated marginal means for constant error model**

Estimated marginal means (EMMs), expressing bias, were computed for all factor combinations.

P-values were adjusted using false discovery rate.

| term | levels | Estimate | Std. Error | df | t value | lower | upper | Pr(>\|t\|) |
|---|---|---|---|---|---|---|---|---|
| posture:direction:region:search | straight:proximodistal:proximal:initial | -0.149 | 0.624 | 1085 | -0.239 | -1.373 | 1.07 | 0.8112 |
| posture:direction:region:search | angled:proximodistal:proximal:initial | 2.523 | 0.640 | 1107 | 3.939 | 1.266 | 3.78 | <0.001 |
| posture:direction:region:search | straight:mediolateral:proximal:initial | -0.574 | 0.624 | 1085 | -0.920 | -1.798 | 0.65 | 0.3579 |
| posture:direction:region:search | angled:mediolateral:proximal:initial | 1.073 | 0.640 | 1107 | 1.676 | -0.183 | 2.33 | 0.0941 |
| posture:direction:region:search | straight:proximodistal:medial:initial | 1.458 | 0.624 | 1085 | 2.337 | 0.234 | 2.68 | 0.0196 |
| posture:direction:region:search | angled:proximodistal:medial:initial | 2.295 | 0.635 | 1099 | 3.615 | 1.049 | 3.54 | <0.001 |
| posture:direction:region:search | straight:mediolateral:medial:initial | 0.545 | 0.624 | 1085 | 0.873 | -0.680 | 1.77 | 0.383 |
| posture:direction:region:search | angled:mediolateral:medial:initial | 2.638 | 0.635 | 1099 | 4.156 | 1.393 | 3.88 | <0.001 |
| posture:direction:region:search | straight:proximodistal:distal:initial | 1.950 | 0.624 | 1085 | 3.126 | 0.726 | 3.17 | 0.0018 |
| posture:direction:region:search | angled:proximodistal:distal:initial | 0.979 | 0.629 | 1092 | 1.556 | -0.256 | 2.21 | 0.1201 |
| posture:direction:region:search | straight:mediolateral:distal:initial | 2.040 | 0.624 | 1085 | 3.270 | 0.816 | 3.27 | 0.0011 |
| posture:direction:region:search | angled:mediolateral:distal:initial | 2.151 | 0.629 | 1092 | 3.418 | 0.916 | 3.39 | <0.001 |
| posture:direction:region:search | straight:proximodistal:proximal:final | 1.157 | 0.624 | 1085 | 1.854 | -0.068 | 2.38 | 0.064 |
| posture:direction:region:search | angled:proximodistal:proximal:final | 0.912 | 0.640 | 1107 | 1.424 | -0.345 | 2.17 | 0.1547 |
| posture:direction:region:search | straight:mediolateral:proximal:final | 0.585 | 0.624 | 1085 | 0.937 | -0.640 | 1.81 | 0.3491 |
| posture:direction:region:search | angled:mediolateral:proximal:final | -0.221 | 0.640 | 1107 | -0.344 | -1.477 | 1.04 | 0.7306 |
| posture:direction:region:search | straight:proximodistal:medial:final | 1.472 | 0.624 | 1085 | 2.359 | 0.248 | 2.70 | 0.0185 |
| posture:direction:region:search | angled:proximodistal:medial:final | 1.141 | 0.635 | 1099 | 1.797 | -0.105 | 2.39 | 0.0726 |
| posture:direction:region:search | straight:mediolateral:medial:final | 1.604 | 0.624 | 1085 | 2.571 | 0.380 | 2.83 | 0.0103 |
| posture:direction:region:search | angled:mediolateral:medial:final | 1.610 | 0.635 | 1099 | 2.537 | 0.365 | 2.86 | 0.0113 |
| posture:direction:region:search | straight:proximodistal:distal:final | 2.750 | 0.624 | 1085 | 4.407 | 1.525 | 3.97 | <0.001 |
| posture:direction:region:search | angled:proximodistal:distal:final | -0.163 | 0.629 | 1092 | -0.259 | -1.398 | 1.07 | 0.7956 |
| posture:direction:region:search | straight:mediolateral:distal:final | 2.443 | 0.624 | 1085 | 3.915 | 1.218 | 3.67 | <0.001 |
| posture:direction:region:search | angled:mediolateral:distal:final | 2.146 | 0.629 | 1092 | 3.410 | 0.911 | 3.38 | <0.001 |

**Table S3: Post-hoc comparisons between target regions (proximal, medial, distal) in different postures (straight vs. angled)**

P-values were adjusted using false discovery rate.

| term | levels | Estimate | Std. Error | df | t value | lower | upper | Pr(>\|t\|) |
|---|---|---|---|---|---|---|---|---|
| posture:region | straight:proximal - angled:proximal | -0.81717 | 0.428 | 1255 | -1.9096 | -1.657 | 0.022 | 0.0564 |
| posture:region | straight:proximal - straight:medial | -1.01515 | 0.420 | 1239 | -2.4195 | -1.838 | -0.192 | 0.0157 |
| posture:region | straight:proximal - angled:medial | -1.66647 | 0.426 | 1254 | -3.9155 | -2.501 | -0.831 | <0.001 |
| posture:region | straight:proximal - straight:distal | -2.04114 | 0.420 | 1239 | -4.8648 | -2.864 | -1.218 | <0.001 |
| posture:region | straight:proximal - angled:distal | -1.02381 | 0.423 | 1252 | -2.4186 | -1.854 | -0.193 | 0.0157 |
| posture:region | angled:proximal - straight:medial | -0.19798 | 0.428 | 1255 | -0.4626 | -1.038 | 0.642 | 0.6437 |
| posture:region | angled:proximal - angled:medial | -0.84929 | 0.430 | 1240 | -1.9771 | -1.692 | -0.007 | 0.0482 |
| posture:region | angled:proximal - straight:distal | -1.22397 | 0.428 | 1255 | -2.8602 | -2.064 | -0.384 | 0.0043 |
| posture:region | angled:proximal - angled:distal | -0.20664 | 0.428 | 1242 | -0.4829 | -1.046 | 0.633 | 0.6293 |
| posture:region | straight:medial - angled:medial | -0.65132 | 0.426 | 1254 | -1.5303 | -1.486 | 0.184 | 0.1262 |
| posture:region | straight:medial - straight:distal | -1.02599 | 0.420 | 1239 | -2.4453 | -1.849 | -0.203 | 0.0146 |
| posture:region | straight:medial - angled:distal | -0.00866 | 0.423 | 1252 | -0.0205 | -0.839 | 0.822 | 0.9837 |
| posture:region | angled:medial - straight:distal | -0.37467 | 0.426 | 1254 | -0.8803 | -1.210 | 0.460 | 0.3788 |
| posture:region | angled:medial - angled:distal | 0.64266 | 0.426 | 1241 | 1.5099 | -0.192 | 1.478 | 0.1313 |
| posture:region | straight:distal - angled:distal | 1.01733 | 0.423 | 1252 | 2.4033 | 0.187 | 1.848 | 0.0164 |

**Table S4: Post-hoc comparisons between directions (proximodistal vs. mediolateral) and search (initial vs. final)**

P-values were adjusted using false discovery rate.

| term | levels | Estimate | Std. Error | df | t value | lower | upper | Pr(>\|t\|) |
|---|---|---|---|---|---|---|---|---|
| direction:search | proximodistal:initial - mediolateral:initial | 0.1970 | 0.346 | 1239 | 0.570 | -0.481 | 0.875 | 0.5689 |
| direction:search | proximodistal:initial - proximodistal:final | 0.2980 | 0.346 | 1239 | 0.862 | -0.380 | 0.976 | 0.3889 |
| direction:search | proximodistal:initial - mediolateral:final | 0.1481 | 0.346 | 1239 | 0.428 | -0.530 | 0.826 | 0.6684 |
| direction:search | mediolateral:initial - proximodistal:final | 0.1010 | 0.346 | 1239 | 0.292 | -0.577 | 0.779 | 0.7703 |
| direction:search | mediolateral:initial - mediolateral:final | -0.0489 | 0.346 | 1239 | -0.141 | -0.727 | 0.629 | 0.8875 |
| direction:search | proximodistal:final - mediolateral:final | -0.1499 | 0.346 | 1239 | -0.433 | -0.828 | 0.528 | 0.6648 |

##### 1.1.2. Mixed model for ansolute (unsigned) error

The factors used in this model are the same as before. However, the predicted variable is the unsigned error, expressing absolute distance from the target.

The resulting formula was:

```
lmer(abs(constError) ~ posture*direction*region*search + (1|id))
```

**Table S5: Anova table for absolute error model**

| Term | Sum Sq | Mean Sq | NumDF | DenDF | F value | Pr(>F) |
|---|---|---|---|---|---|---|
| posture | 0.407 | 0.407 | 1 | 1273 | 0.047 | 0.8283 |
| direction | 833.139 | 833.139 | 1 | 1240 | 96.328 | <0.001 |
| region | 32.297 | 16.149 | 2 | 1241 | 1.867 | 0.155 |
| search | 165.879 | 165.879 | 1 | 1240 | 19.179 | <0.001 |
| posture:direction | 23.833 | 23.833 | 1 | 1240 | 2.756 | 0.0972 |
| posture:region | 120.245 | 60.122 | 2 | 1241 | 6.951 | 0.001 |
| direction:region | 35.647 | 17.823 | 2 | 1240 | 2.061 | 0.1278 |
| posture:search | 31.692 | 31.692 | 1 | 1240 | 3.664 | 0.0558 |
| direction:search | 4.013 | 4.013 | 1 | 1240 | 0.464 | 0.4959 |
| region:search | 17.349 | 8.674 | 2 | 1240 | 1.003 | 0.3671 |
| posture:direction:region | 59.049 | 29.525 | 2 | 1240 | 3.414 | 0.0332 |
| posture:direction:search | 5.466 | 5.466 | 1 | 1240 | 0.632 | 0.4268 |
| posture:region:search | 3.160 | 1.580 | 2 | 1240 | 0.183 | 0.8331 |
| direction:region:search | 8.494 | 4.247 | 2 | 1240 | 0.491 | 0.6121 |
| posture:direction:region:search | 9.358 | 4.679 | 2 | 1240 | 0.541 | 0.5823 |

**Table S6: Post-hoc comparisons between directions (proximodistal vs. mediolateral) and search (initial vs. final)**

P-values were adjusted using false discovery rate.

| term | levels | Estimate | Std. Error | df | t value | lower | upper | Pr(>\|t\|) |
|---|---|---|---|---|---|---|---|---|
| direction:search | proximodistal:initial - mediolateral:initial | 1.700 | 0.229 | 1240 | 7.42 | 1.25 | 2.149 | <0.001 |
| direction:search | proximodistal:initial - proximodistal:final | 0.819 | 0.229 | 1240 | 3.58 | 0.37 | 1.269 | <0.001 |
| direction:search | proximodistal:initial - mediolateral:final | 2.298 | 0.229 | 1240 | 10.04 | 1.85 | 2.748 | <0.001 |
| direction:search | mediolateral:initial - proximodistal:final | -0.880 | 0.229 | 1240 | -3.84 | -1.33 | -0.431 | <0.001 |
| direction:search | mediolateral:initial - mediolateral:final | 0.599 | 0.229 | 1240 | 2.62 | 0.15 | 1.048 | 0.009 |
| direction:search | proximodistal:final - mediolateral:final | 1.479 | 0.229 | 1240 | 6.46 | 1.03 | 1.928 | <0.001 |

###### 1.1.3. Linear mixed model for variable error

The same factors were used for the variable error model. The resulting formula was:

```
lmer(varError ~ posture*direction*region*search + (1|id))
```

**Table S7: Anova table for variable error model**

| Term | Sum Sq | Mean Sq | NumDF | DenDF | F value | Pr(>F) |
|---|---|---|---|---|---|---|
| posture | 22.60558 | 22.60558 | 1 | 1261 | 5.371 | 0.0206 |
| direction | 1389.42223 | 1389.42223 | 1 | 1223 | 330.129 | <0.001 |
| region | 293.35628 | 146.67814 | 2 | 1225 | 34.851 | <0.001 |
| search | 733.74641 | 733.74641 | 1 | 1223 | 174.339 | <0.001 |
| posture:direction | 1.83419 | 1.83419 | 1 | 1223 | 0.436 | 0.5093 |
| posture:region | 121.52490 | 60.76245 | 2 | 1225 | 14.437 | <0.001 |
| direction:region | 106.89537 | 53.44768 | 2 | 1223 | 12.699 | <0.001 |
| posture:search | 1.48640 | 1.48640 | 1 | 1223 | 0.353 | 0.5524 |
| direction:search | 0.00902 | 0.00902 | 1 | 1223 | 0.002 | 0.9631 |
| region:search | 13.70274 | 6.85137 | 2 | 1223 | 1.628 | 0.1968 |
| posture:direction:region | 37.57302 | 18.78651 | 2 | 1223 | 4.464 | 0.0117 |
| posture:direction:search | 0.01060 | 0.01060 | 1 | 1223 | 0.003 | 0.96 |
| posture:region:search | 14.17559 | 7.08780 | 2 | 1223 | 1.684 | 0.186 |
| direction:region:search | 3.72341 | 1.86171 | 2 | 1223 | 0.442 | 0.6426 |
| posture:direction:region:search | 4.51546 | 2.25773 | 2 | 1223 | 0.536 | 0.585 |

**Table S8: Post-hoc comparison between target regions (proximal, medial, distal)**

P-values were adjusted using false discovery rate.

| term | levels | Estimate | Std. Error | df | t value | lower | upper | Pr(>\|t\|) |
|---|---|---|---|---|---|---|---|---|
| region | proximal - medial | -0.112 | 0.139 | 1226 | -0.802 | -0.386 | 0.162 | 0.4227 |
| region | proximal - distal | 0.945 | 0.140 | 1227 | 6.757 | 0.670 | 1.219 | <0.001 |
| region | medial - distal | 1.057 | 0.139 | 1223 | 7.619 | 0.785 | 1.329 | <0.001 |

**Table S9: Post-hoc comparison between directions (proximodistal vs. mediolateral)**

P-values were adjusted using false discovery rate.

| term | levels | Estimate | Std. Error | df | t value | lower | upper | Pr(>\|t\|) |
|---|---|---|---|---|---|---|---|---|
| direction | proximodistal - mediolateral | 2.07 | 0.114 | 1223 | 18.2 | 1.84 | 2.29 | <0.001 |

**Table S10: Post-hoc comparisons between search (initial vs. final) for postures (straight vs. angled)**

P-values were adjusted using false discovery rate.

| term | levels | Estimate | Std. Error | df | t value | lower | upper | Pr(>\|t\|) |
|---|---|---|---|---|---|---|---|---|
| posture:search | straight:initial - angled:initial | -0.200 | 0.162 | 1244 | -1.23 | -0.518 | 0.118 | 0.2173 |
| posture:search | straight:initial - straight:final | 1.568 | 0.159 | 1223 | 9.85 | 1.256 | 1.881 | <0.001 |
| posture:search | straight:initial - angled:final | 1.233 | 0.162 | 1244 | 7.61 | 0.915 | 1.551 | <0.001 |

| term | levels | Estimate | Std. Error | df | t value | lower | upper | Pr(>\|t\|) |
|---|---|---|---|---|---|---|---|---|
| posture:search | angled:initial - straight:final | 1.768 | 0.162 | 1244 | 10.92 | 1.450 | 2.086 | <0.001 |
| posture:search | angled:initial - angled:final | 1.433 | 0.162 | 1223 | 8.83 | 1.115 | 1.751 | <0.001 |
| posture:search | straight:final - angled:final | -0.335 | 0.162 | 1244 | -2.07 | -0.653 | -0.017 | 0.0388 |

#### 1.2. Characterization of error reduction during search

##### 1.2.1. Linear mixed model for search model

With the following models, we analyzed the effect of the travelled distance (percent moved) during the (spatialized) trajectories on direction-independent localization error (computed as Euclidian distance from the target).

The full model contained the following factors:

- fixed factors:
  - posture (straight, angled)
  - target region (proximal, medial, distal)
  - initial localization error (small, medium, large)
  - linear othogocal polynomial term for percent moved (coded in 21 segments representing 5% moved)
  - quadratic othogonal polynomial term for percent moved
- random factors:
  - id (random intercept factor)

The resulting model formula was:

```
lmer(tdists_percent_arm ~ posture*region*initialError*percent_moved.L*percent_moved.Q +
(1|id))
```

**Table S11: Anova table for search model**

| Term | Sum Sq | Mean Sq | NumDF | DenDF | F value | Pr(>F) |
|---|---|---|---|---|---|---|
| posture | 28.04 | 28.045 | 1 | 31007 | 2.790 | 0.0948 |
| region | 1192.06 | 596.030 | 2 | 30958 | 59.303 | <0.001 |
| initialError | 168180.69 | 84090.344 | 2 | 31006 | 8366.672 | <0.001 |
| percent__moved.L | 3911.48 | 3911.479 | 1 | 30951 | 389.177 | <0.001 |
| percent__moved.Q | 772.27 | 772.275 | 1 | 30951 | 76.838 | <0.001 |
| posture:region | 4743.87 | 2371.935 | 2 | 30960 | 235.999 | <0.001 |
| posture:initialError | 950.23 | 475.114 | 2 | 30989 | 47.272 | <0.001 |
| region:initialError | 3785.20 | 946.299 | 4 | 30985 | 94.153 | <0.001 |
| posture:percent__moved.L | 5.55 | 5.545 | 1 | 30951 | 0.552 | 0.4576 |
| region:percent__moved.L | 122.09 | 61.044 | 2 | 30951 | 6.074 | 0.0023 |
| initialError:percent__moved.L | 7825.79 | 3912.893 | 2 | 30951 | 389.318 | <0.001 |
| posture:percent__moved.Q | 111.29 | 111.294 | 1 | 30951 | 11.073 | <0.001 |
| region:percent__moved.Q | 46.20 | 23.099 | 2 | 30951 | 2.298 | 0.1005 |
| initialError:percent__moved.Q | 2788.18 | 1394.090 | 2 | 30951 | 138.707 | <0.001 |
| percent__moved.L:percent__moved.Q | 104.35 | 104.354 | 1 | 30951 | 10.383 | 0.0013 |
| posture:region:initialError | 1212.79 | 303.197 | 4 | 30983 | 30.167 | <0.001 |
| posture:region:percent__moved.L | 73.15 | 36.577 | 2 | 30951 | 3.639 | 0.0263 |
| posture:initialError:percent__moved.L | 30.70 | 15.352 | 2 | 30951 | 1.527 | 0.2171 |
| region:initialError:percent__moved.L | 292.81 | 73.203 | 4 | 30951 | 7.283 | <0.001 |
| posture:region:percent__moved.Q | 158.48 | 79.239 | 2 | 30951 | 7.884 | <0.001 |
| posture:initialError:percent__moved.Q | 43.28 | 21.642 | 2 | 30951 | 2.153 | 0.1161 |
| region:initialError:percent__moved.Q | 79.93 | 19.982 | 4 | 30951 | 1.988 | 0.0934 |
| posture:percent__moved.L:percent__moved.Q | 15.06 | 15.063 | 1 | 30951 | 1.499 | 0.2209 |
| region:percent__moved.L:percent__moved.Q | 4.34 | 2.169 | 2 | 30951 | 0.216 | 0.8059 |
| initialError:percent__moved.L:percent__moved.Q | 13.02 | 6.512 | 2 | 30951 | 0.648 | 0.5231 |
| posture:region:initialError:percent__moved.L | 117.76 | 29.441 | 4 | 30951 | 2.929 | 0.0196 |
| posture:region:initialError:percent__moved.Q | 18.91 | 4.728 | 4 | 30951 | 0.470 | 0.7575 |
| posture:region:percent__moved.L:percent__moved.Q | 1.10 | 0.552 | 2 | 30951 | 0.055 | 0.9465 |
| posture:initialError:percent__moved.L:percent__moved.Q | 3.50 | 1.750 | 2 | 30951 | 0.174 | 0.8402 |
| region:initialError:percent__moved.L:percent__moved.Q | 52.63 | 13.158 | 4 | 30951 | 1.309 | 0.2639 |
| posture:region:initialError:percent__moved.L:percent__moved.Q | 29.24 | 7.309 | 4 | 30951 | 0.727 | 0.5732 |

##### 1.2.2. Linear and quaratic terms for different initial localization error

To compare the different slopes in error reduction through searh for different initial distances, we computed a reduced model, containing only initial distance and the linear and quadratic terms of percent moved.

The mdeol forlula was:

```
lmer(tdists_percent_arm ~ initialError+percent_moved.L + percent_moved.Q + initialError:percent_moved.L
      + initialError:percent_moved.Q + (1|id)
```

**Table S12: Regression table for the reduced search model**

| Term | Estimate | Std. Error | df | t value | Pr(>\|t\|) |
|---|---|---|---|---|---|
| (Intercept) | 4.530 | 0.1501 | 60.8 | 30.18 | <0.001 |
| initialErrormedium | 2.228 | 0.0461 | 31051.9 | 48.38 | <0.001 |
| initialErrorlarge | 6.396 | 0.0480 | 31064.1 | 133.20 | <0.001 |
| percent_moved.L | 0.834 | 0.1462 | 31013.8 | 5.70 | <0.001 |
| percent_moved.Q | -0.927 | 0.1462 | 31013.8 | -6.34 | <0.001 |
| initialErrormedium:percent_moved.L | -2.628 | 0.2069 | 31013.8 | -12.70 | <0.001 |
| initialErrorlarge:percent_moved.L | -7.823 | 0.2054 | 31013.8 | -38.09 | <0.001 |
| initialErrormedium:percent_moved.Q | 1.660 | 0.2069 | 31013.8 | 8.02 | <0.001 |
| initialErrorlarge:percent_moved.Q | 3.325 | 0.2054 | 31013.8 | 16.19 | <0.001 |

Note that the regression coefficients in the table are expressed as treatment contrasts (the default in R). That means, the intercept expresses the estimation for the combination of the baseline level for all factors. The baseline level for the initial localization error is "small".

- The terms for the linear component therefore compute as:
  - initial small error (baseline): 0.834
  - initial medium error: 0.834 -2.628 = -1.794
  - initial large error: 0.834 -7.823 = -6.989
- The terms for the quadratic component therefore compute as:
  - initial small error (baseline): -0.927
  - initial medium error: -0.927 + 1.66 = 0.733
  - initial large error: -0.927 + 3.325 = 2.398

**Table S13: Linear and quadratic terms for different initial error (small, medium, large), postures (straight vs. angled) and target regions (proximal, medial, distal)**

To test whether the components were significant for the combinations of initial postures and target regions, the model was repeadedly fit in subsets of the data. The results are summarized in Table S13.

| | initialError | posture | region | Term | Estimate | Std. Error | df | t value | Pr(>\|t\|) |
|---|---|---|---|---|---|---|---|---|---|
| 2 | small | angled | distal | percent_moved.L | 0.0918 | 0.160 | 2175 | 0.573 | 0.5664 |
| 21 | small | angled | medial | percent_moved.L | 0.7629 | 0.206 | 1469 | 3.710 | <0.001 |
| 22 | small | angled | proximal | percent_moved.L | 1.1038 | 0.224 | 1531 | 4.919 | <0.001 |
| 23 | small | straight | distal | percent_moved.L | 2.1031 | 0.229 | 1491 | 9.190 | <0.001 |
| 24 | small | straight | medial | percent_moved.L | 0.8727 | 0.191 | 1511 | 4.558 | <0.001 |
| 25 | small | straight | proximal | percent_moved.L | 0.4747 | 0.189 | 1826 | 2.514 | 0.012 |
| 26 | medium | angled | distal | percent_moved.L | -2.0416 | 0.223 | 1922 | -9.163 | <0.001 |
| 27 | medium | angled | medial | percent_moved.L | -1.5555 | 0.236 | 1633 | -6.598 | <0.001 |
| 28 | medium | angled | proximal | percent_moved.L | -1.4791 | 0.278 | 1241 | -5.315 | <0.001 |
| 29 | medium | straight | distal | percent_moved.L | -2.5121 | 0.193 | 1675 | -13.022 | <0.001 |

| | initialError | posture | region | Term | Estimate | Std. Error | df | t value | Pr(>\|t\|) |
|---|---|---|---|---|---|---|---|---|---|
| 210 | medium | straight | medial | percent_moved.L | -1.1035 | 0.205 | 1591 | -5.380 | <0.001 |
| 211 | medium | straight | proximal | percent_moved.L | -1.9027 | 0.219 | 1905 | -8.688 | <0.001 |
| 212 | large | angled | distal | percent_moved.L | -7.8636 | 0.416 | 1099 | -18.881 | <0.001 |
| 213 | large | angled | medial | percent_moved.L | -6.1742 | 0.359 | 1926 | -17.207 | <0.001 |
| 214 | large | angled | proximal | percent_moved.L | -8.3330 | 0.376 | 2156 | -22.154 | <0.001 |
| 215 | large | straight | distal | percent_moved.L | -6.2961 | 0.287 | 1509 | -21.933 | <0.001 |
| 216 | large | straight | medial | percent_moved.L | -5.1308 | 0.298 | 1988 | -17.206 | <0.001 |
| 217 | large | straight | proximal | percent_moved.L | -8.5049 | 0.393 | 1620 | -21.657 | <0.001 |
| 3 | small | angled | distal | percent_moved.Q | -0.4547 | 0.160 | 2175 | -2.839 | 0.0046 |
| 31 | small | angled | medial | percent_moved.Q | -0.7793 | 0.206 | 1469 | -3.790 | <0.001 |
| 32 | small | angled | proximal | percent_moved.Q | -1.1312 | 0.224 | 1531 | -5.041 | <0.001 |
| 33 | small | straight | distal | percent_moved.Q | -1.9486 | 0.229 | 1491 | -8.515 | <0.001 |
| 34 | small | straight | medial | percent_moved.Q | -0.8716 | 0.191 | 1511 | -4.553 | <0.001 |
| 35 | small | straight | proximal | percent_moved.Q | -0.6473 | 0.189 | 1826 | -3.429 | <0.001 |
| 36 | medium | angled | distal | percent_moved.Q | 0.9737 | 0.223 | 1922 | 4.370 | <0.001 |
| 37 | medium | angled | medial | percent_moved.Q | 0.7353 | 0.236 | 1633 | 3.119 | 0.0018 |
| 38 | medium | angled | proximal | percent_moved.Q | 0.8772 | 0.278 | 1241 | 3.152 | 0.0017 |
| 39 | medium | straight | distal | percent_moved.Q | 0.1552 | 0.193 | 1675 | 0.805 | 0.4211 |
| 310 | medium | straight | medial | percent_moved.Q | 0.8546 | 0.205 | 1591 | 4.166 | <0.001 |
| 311 | medium | straight | proximal | percent_moved.Q | 0.7994 | 0.219 | 1905 | 3.650 | <0.001 |
| 312 | large | angled | distal | percent_moved.Q | 3.5118 | 0.416 | 1099 | 8.432 | <0.001 |
| 313 | large | angled | medial | percent_moved.Q | 2.1935 | 0.359 | 1926 | 6.113 | <0.001 |
| 314 | large | angled | proximal | percent_moved.Q | 3.1782 | 0.376 | 2156 | 8.449 | <0.001 |
| 315 | large | straight | distal | percent_moved.Q | 1.3351 | 0.287 | 1509 | 4.651 | <0.001 |
| 316 | large | straight | medial | percent_moved.Q | 1.5797 | 0.298 | 1988 | 5.298 | <0.001 |
| 317 | large | straight | proximal | percent_moved.Q | 2.8376 | 0.393 | 1620 | 7.225 | <0.001 |

### 2. Experiment 2

#### 2.1. Statistical models for constant and variable error

##### 2.1.1. Linear mixed model for constant error

The following factors were included in the model:

- fixed factors:
  - surface (skin, barrier)
  - direction (proximodistal, mediolateral)
  - search (initial final)
- random factors:
  - id (random intercept factor)

resulting formula in lme4 notation: `lmer(constError ~ surface*direction*search + (1|id))`

**Table S14: Anova table for constant error model**

| Term | Sum Sq | Mean Sq | NumDF | DenDF | F value | Pr(>F) |
|---|---|---|---|---|---|---|
| surface | 572.46 | 572.46 | 1 | 233 | 10.704 | 0.0012 |
| direction | 4869.65 | 4869.65 | 1 | 233 | 91.052 | <0.001 |
| search | 322.42 | 322.42 | 1 | 233 | 6.029 | 0.0148 |

| Term | Sum Sq | Mean Sq | NumDF | DenDF | F value | Pr(>F) |
|---|---|---|---|---|---|---|
| surface:direction | 5558.87 | 5558.87 | 1 | 233 | 103.939 | <0.001 |
| surface:search | 31.13 | 31.13 | 1 | 233 | 0.582 | 0.4463 |
| direction:search | 0.55 | 0.55 | 1 | 233 | 0.010 | 0.9193 |
| surface:direction:search | 232.25 | 232.25 | 1 | 233 | 4.343 | 0.0383 |

**Table S15: Estimated marginal means for constant error model**

Estimated marginal means (EMMs), expressing bias, were computed for all factor combinations.

P-values were adjusted using false discovery rate.

| term | levels | Estimate | Std. Error | df | t value | lower | upper | Pr(>\|t\|) |
|---|---|---|---|---|---|---|---|---|
| surface:direction:search | skin:proximodistal:initial | -0.784 | 1.55 | 85.6 | -0.507 | -3.86 | 2.291 | 0.6134 |
| surface:direction:search | barrier:proximodistal:initial | -4.511 | 1.55 | 85.6 | -2.917 | -7.59 | -1.436 | 0.0045 |
| surface:direction:search | skin:mediolateral:initial | 0.431 | 1.55 | 85.6 | 0.279 | -2.64 | 3.506 | 0.7811 |
| surface:direction:search | barrier:mediolateral:initial | 11.534 | 1.55 | 85.6 | 7.457 | 8.46 | 14.609 | <0.001 |
| surface:direction:search | skin:proximodistal:final | -0.519 | 1.55 | 85.6 | -0.336 | -3.59 | 2.556 | 0.738 |
| surface:direction:search | barrier:proximodistal:final | -9.451 | 1.55 | 85.6 | -6.110 | -12.53 | -6.376 | <0.001 |
| surface:direction:search | skin:mediolateral:final | -2.928 | 1.55 | 85.6 | -1.893 | -6.00 | 0.147 | 0.0617 |
| surface:direction:search | barrier:mediolateral:final | 10.590 | 1.55 | 85.6 | 6.847 | 7.51 | 13.665 | <0.001 |

##### 2.1.2. Linear mixed model for variable error

The same factors were used for the variable error model. The resulting formula was:

```
lmer(varError ~ surface*direction*search + (1|id))
```

**Table S16: Anova table for variable error model**

| Term | Sum Sq | Mean Sq | NumDF | DenDF | F value | Pr(>F) |
|---|---|---|---|---|---|---|
| surface | 1355.98 | 1355.98 | 1 | 233 | 131.863 | <0.001 |
| direction | 567.69 | 567.69 | 1 | 233 | 55.205 | <0.001 |
| search | 227.48 | 227.48 | 1 | 233 | 22.122 | <0.001 |
| surface:direction | 148.84 | 148.84 | 1 | 233 | 14.474 | <0.001 |
| surface:search | 33.75 | 33.75 | 1 | 233 | 3.282 | 0.0713 |
| direction:search | 4.73 | 4.73 | 1 | 233 | 0.460 | 0.4983 |
| surface:direction:search | 2.59 | 2.59 | 1 | 233 | 0.252 | 0.616 |

#### 2.3. Characterization of error reduction during search

##### 2.3.1. Linear mixed model for search

The model contained the following factors:

- fixed factors:
  - surface (skin, bariier)
  - initial localization error (small, medium, large)
  - linear othogocal polynomial term for percent moved (coded in 21 segments representing 5% moved)
  - quadratic othogonal polynomial term for percent moved

- random factors:
  - id (random intercept factor)

The resulting model formula was:

```
lmer(tdists_percent_arm ~ surface*percent_moved.L*percent_moved.Q + (1|id))
```

**Table S17: Anova table for the search model**

| Term | Sum Sq | Mean Sq | NumDF | DenDF | F value | Pr(>F) |
|---|---|---|---|---|---|---|
| surface | 58635.297 | 58635.297 | 1 | 6087 | 1671.190 | <0.001 |
| initialError | 61376.207 | 30688.103 | 2 | 6041 | 874.655 | <0.001 |
| percent_moved.L | 432.347 | 432.347 | 1 | 6072 | 12.323 | <0.001 |
| percent_moved.Q | 1011.260 | 1011.260 | 1 | 6072 | 28.822 | <0.001 |
| surface:initialError | 20954.397 | 10477.198 | 2 | 6086 | 298.615 | <0.001 |
| surface:percent_moved.L | 3766.320 | 3766.320 | 1 | 6072 | 107.346 | <0.001 |
| initialError:percent_moved.L | 5166.004 | 2583.002 | 2 | 6072 | 73.619 | <0.001 |
| surface:percent_moved.Q | 668.494 | 668.494 | 1 | 6072 | 19.053 | <0.001 |
| initialError:percent_moved.Q | 1365.307 | 682.654 | 2 | 6072 | 19.457 | <0.001 |
| percent_moved.L:percent_moved.Q | 198.456 | 198.456 | 1 | 6072 | 5.656 | 0.0174 |
| surface:initialError:percent_moved.L | 391.293 | 195.646 | 2 | 6072 | 5.576 | 0.0038 |
| surface:initialError:percent_moved.Q | 1589.935 | 794.967 | 2 | 6072 | 22.658 | <0.001 |
| surface:percent_moved.L:percent_moved.Q | 1.128 | 1.128 | 1 | 6072 | 0.032 | 0.8577 |
| initialError:percent_moved.L:percent_moved.Q | 0.022 | 0.011 | 2 | 6072 | 0.000 | 0.9997 |
| surface:initialError:percent_moved.L:percent_moved.Q | 164.767 | 82.383 | 2 | 6072 | 2.348 | 0.0956 |

**Table S18: Linear and quadratic terms for different initial error (small, medium, large) and surfaces (skin vs. barrier)**

To test whether the components were significant for the combinations of initial localization error and and surfaces, the model was repeadedly fit in subsets of the data. The results are summarized in Table S18.

| | initialError | surface | Term | Estimate | Std. Error | df | t value | Pr(>\|t\|) |
|---|---|---|---|---|---|---|---|---|
| 2 | small | barrier | percent_moved.L | 8.5172 | 0.759 | 411 | 11.2273 | <0.001 |
| 21 | small | skin | percent_moved.L | 0.9775 | 0.314 | 1579 | 3.1101 | 0.0019 |
| 22 | medium | barrier | percent_moved.L | 3.3072 | 0.666 | 950 | 4.9694 | <0.001 |
| 23 | medium | skin | percent_moved.L | -5.7917 | 0.440 | 1033 | -13.1601 | <0.001 |
| 24 | large | barrier | percent_moved.L | -3.3532 | 0.988 | 1536 | -3.3930 | <0.001 |
| 25 | large | skin | percent_moved.L | -21.2460 | 0.838 | 515 | -25.3561 | <0.001 |
| 26 | all trials | barrier | percent_moved.L | 0.5310 | 0.707 | 2922 | 0.7507 | 0.4529 |
| 27 | all trials | skin | percent_moved.L | -4.9433 | 0.320 | 3153 | -15.4441 | <0.001 |
| 3 | small | barrier | percent_moved.Q | 0.0526 | 0.759 | 411 | 0.0693 | 0.9448 |
| 31 | small | skin | percent_moved.Q | -1.8535 | 0.314 | 1579 | -5.8972 | <0.001 |
| 32 | medium | barrier | percent_moved.Q | 1.2044 | 0.666 | 950 | 1.8098 | 0.0706 |
| 33 | medium | skin | percent_moved.Q | 2.5337 | 0.440 | 1033 | 5.7572 | <0.001 |
| 34 | large | barrier | percent_moved.Q | -0.0739 | 0.988 | 1536 | -0.0748 | 0.9404 |
| 35 | large | skin | percent_moved.Q | 10.7935 | 0.838 | 515 | 12.8815 | <0.001 |
| 36 | all trials | barrier | percent_moved.Q | 0.3642 | 0.707 | 2922 | 0.5149 | 0.6067 |
| 37 | all trials | skin | percent_moved.Q | 1.6931 | 0.320 | 3153 | 5.2896 | <0.001 |
